# Cohesin and microtubule dependent mechanisms regulate sister centromere fusion during meiosis I

**DOI:** 10.1101/571067

**Authors:** Lin-Ing Wang, Arunika Das, Kim S. McKim

**Affiliations:** Waksman Institute and Department of Genetics, Rutgers, the State University of New Jersey, Piscataway N.J. 08854-8020

**Keywords:** meiosis, cohesion, chromosome segregation, centromere, kinetochore

## Abstract

Sister centromere fusion is a process unique to meiosis that promotes co-orientation of the sister kinetochores, ensuring they attach to microtubules from the same pole. We have found that the kinetochore protein SPC105R/KNL1 and Protein Phosphatase 1 (PP1-87B) are required for this process. The analysis of these two proteins, however, has shown that two independent mechanisms maintain sister centromere fusion during meiosis I in *Drosophila* oocytes. Double depletion experiments demonstrated that the precocious separation of centromeres in *Spc105R* RNAi oocytes is Separase-dependent, suggesting cohesin proteins must be maintained at the core centromeres. In contrast, precocious sister centromere separation in *Pp1-87B* RNAi oocytes does not depend on Separase or Wapl. Further analysis with microtubule destabilizing drugs showed that PP1-87B maintains sister centromeres fusion by regulating microtubule dynamics. Additional double depletion experiments demonstrated that PP1-87B has this function by antagonizing Polo kinase and BubR1, two proteins known to promote kinetochore-microtubule (KT-MT) attachments. These results suggest that PP1-87B maintains sister centromere fusion by inhibiting stable KT-MT attachments. Surprisingly, we found that loss of C(3)G, the transverse element of the synaptonemal complex (SC), suppresses centromere separation in *Pp1-87B* RNAi oocytes. This is evidence for a functional role of centromeric SC in the meiotic divisions. We propose two mechanisms maintain co-orientation in *Drosophila* oocytes: one involves SPC105R to protect cohesins at sister centromeres and another involves PP1-87B to regulate spindle forces at end-on attachments.

**Author Summary:** Meiosis involves two cell divisions. In the first division, pairs of homologous chromosomes segregate, in the second division, the sister chromatids segregate. These patterns of division are mediated by regulating microtubule attachments to the kinetochores and stepwise release of cohesion between the sister chromatids. During meiosis I, cohesion fusing sister centromeres must be intact so they attach to microtubules from the same pole. At the same time, arm cohesion must be released for anaphase I. Upon entry into meiosis II, the sister centromeres must separate to allow attachment to opposite poles, while cohesion surrounding the centromeres must remain intact until anaphase II. How these different populations of cohesion are regulated is not understood. We identified two genes required for maintaining sister centromere cohesion, and surprisingly found they define two distinct mechanisms. The first is a kinetochore protein that maintains sister centromere fusion by recruiting proteins that protect cohesion. The second is a phosphatase that antagonizes proteins that stabilize microtubule attachments. We propose that entry into meiosis II coincides with stabilization of microtubule attachments, resulting in the separation of sister centromeres without disrupting cohesion in other regions, facilitating attachment of sister chromatids to opposite poles.

## Introduction

The necessity of sister kinetochores to co-orient toward the same pole for co-segregation at anaphase I differentiates the first meiotic division from the second division. A meiosis-specific mechanism exists that ensures sister chromatid co-segregation by rearranging sister kinetochores, aligning them next to each other and facilitating microtubule attachments to the same pole [1, 2]. We refer to this process as co-orientation, in contrast to mono-orientation, when homologous kinetochores orient to the same pole. Given the importance of co-orientation in meiosis the mechanism underlying this process is still poorly understood, maybe because many of the essential proteins are not conserved across phyla.

Most studies of co-orientation have focused on how fusion of the centromeres and kinetochores is established. In budding yeast, centromere fusion occurs independently of cohesins: Spo13 and the Polo kinase homolog Cdc5 recruit a meiosis-specific protein complex, monopolin (Csm1, Lrs4, Mam1, CK1) to the kinetochore [3-5]. Lrs4 and Csm1 form a V-shaped structure that interacts with the N-terminal domain of Dsn1 in the Mis12 complex to fuse sister kinetochores [6, 7]. While the monopolin complex is not widely conserved, cohesin-independent mechanisms may exist in other organisms. A bridge between the kinetochore proteins MIS12 and NDC80 is required for co-orientation in maize [8]. In contrast, cohesins are required for co-orientation in several organisms. The meiosis-specific cohesin Rec8 is indispensable for sister centromere fusion in fission yeast [9] and *Arabidopsis* [10, 11]. Cohesin is localized to the core-centromere in fission yeast [12] and mice [13]. In *Drosophila melanogaster* oocytes, we and others have shown that cohesins (SMC1/SMC3/SOLO/SUNN) establish cohesion in meiotic S-phase and are enriched at the core centromeres [14-17]. Like fission yeast and mouse, *Drosophila* may require high concentrations of cohesins to fuse sister centromeres together for co-orientation during meiosis.

In mice, a novel kinetochore protein, Meikin, recruits Plk1 to protect Rec8 at centromeres [13]. Although poorly conserved, Meikin is proposed to be a functional homolog of Spo13 in budding yeast and Moa1 in fission yeast. They all contain Polo-box domains that recruit Polo kinase to centromeres [13]. Loss of Polo in both fission yeast (Plo1) and mice results in kinetochore separation [13, 18], suggesting a conserved role for Polo in co-orientation. In fission yeast, Moa1-Plo1 phosphorylates Spc7 (KNL1) to recruit Bub1 and Sgo1 for the protection of centromere cohesion in meiosis I [18, 19]. These results suggest the mechanism for maintaining sister centromere fusion involves kinetochore proteins recruiting proteins that protect cohesion. However, how centromere cohesion is established prior to metaphase I, and how sister centromere fusion is released during meiosis II, still needs to be investigated.

We previously found that depletion of the kinetochore protein SPC105R (KNL1) in *Drosophila* oocytes results in separated centromeres at metaphase I, suggesting a defect in sister centromere fusion [20]. Thus, *Drosophila* SPC105R and fission yeast Spc7 may have conserved functions in co-orientation [18]. We have identified a second *Drosophila* protein required for sister-centromere fusion, Protein Phosphatase 1 isoform 87B (PP1-87B). However, sister centromere separation in SPC105R and PP1-87B depleted *Drosophila* oocytes occurs by different mechanisms, the former is Separase dependent and the latter is Separase independent. Based on these results, we propose a model for the establishment, protection and release of co-orientation. Sister centromere fusion necessary for co-orientation is established through cohesins that are protected by SPC105R. Subsequently, PP1-87B maintains co-orientation in a cohesin-independent manner by antagonizing stable kinetochore-microtubule (KT-MT) interactions. The implication is that the release of co-orientation during meiosis II is cohesin-independent and MT dependent. We also found a surprising interaction between PP1-87B and C(3)G, the transverse element of the synaptonemal complex (SC), in regulating sister centromere separation. Overall, our results suggest a new mechanism where KT-MT interactions and centromeric SC regulate sister kinetochore co-orientation during female meiosis.

## Results

### PP1-87B is required for karyosome organization sister centromere fusion in meiosis I

*Drosophila* has three homologs of the mammalian Phosphatase1 (PP1α/γ) genes, *Pp1-87B*, *Pp1-96A* and *Pp1-13C*. We focused our studies on the *Pp1-87B* isoform because it is the only essential gene, is highly expressed during oogenesis, and contributes ~80% of PP1 activity during development [21, 22]. As *Pp1-87B* mutations are lethal, tissue-specific expression of an shRNA targeting *Pp1-87B* was used to define its role in oocytes (see Methods) [23]. The ubiquitous expression of an shRNA for PP1-87B using *tubP-GAL4-LL7* resulted in lethality, suggesting the protein had been depleted. When PP1-87B was depleted in oocytes using *mata4-GAL-VP16* (to be referred to as *Pp1-87B* RNAi oocytes), we observed two phenotypes. The first was disorganization of the metaphase I chromosomes. In wild-type *Drosophila* oocytes, meiosis arrests at metaphase I with the chromosomes clustered into a single spherical structure, the karyosome, at the center of the spindle (Figure 1A). In 62% of *Pp1-87B* RNAi oocytes, the karyosome was separated into multiple groups of chromosomes (Figure 1A, B). The second phenotype observed in *Pp1-87B* RNAi oocytes was precocious separation of sister centromeres, as determined by counting the number of centromere protein CENP-C or CID (CENP-A) foci (see Methods) [24]. In wild-type oocytes, we observed an average of 7.3 centromere foci, consistent with the eight expected from four bivalent chromosomes at metaphase I (Figure 1A, C). However, in *Pp1-87B* RNAi oocytes we observed a significantly higher number of foci (mean=12.7). This suggests a defect in sister centromere fusion results in their premature separation during metaphase I.

**Figure 1:**
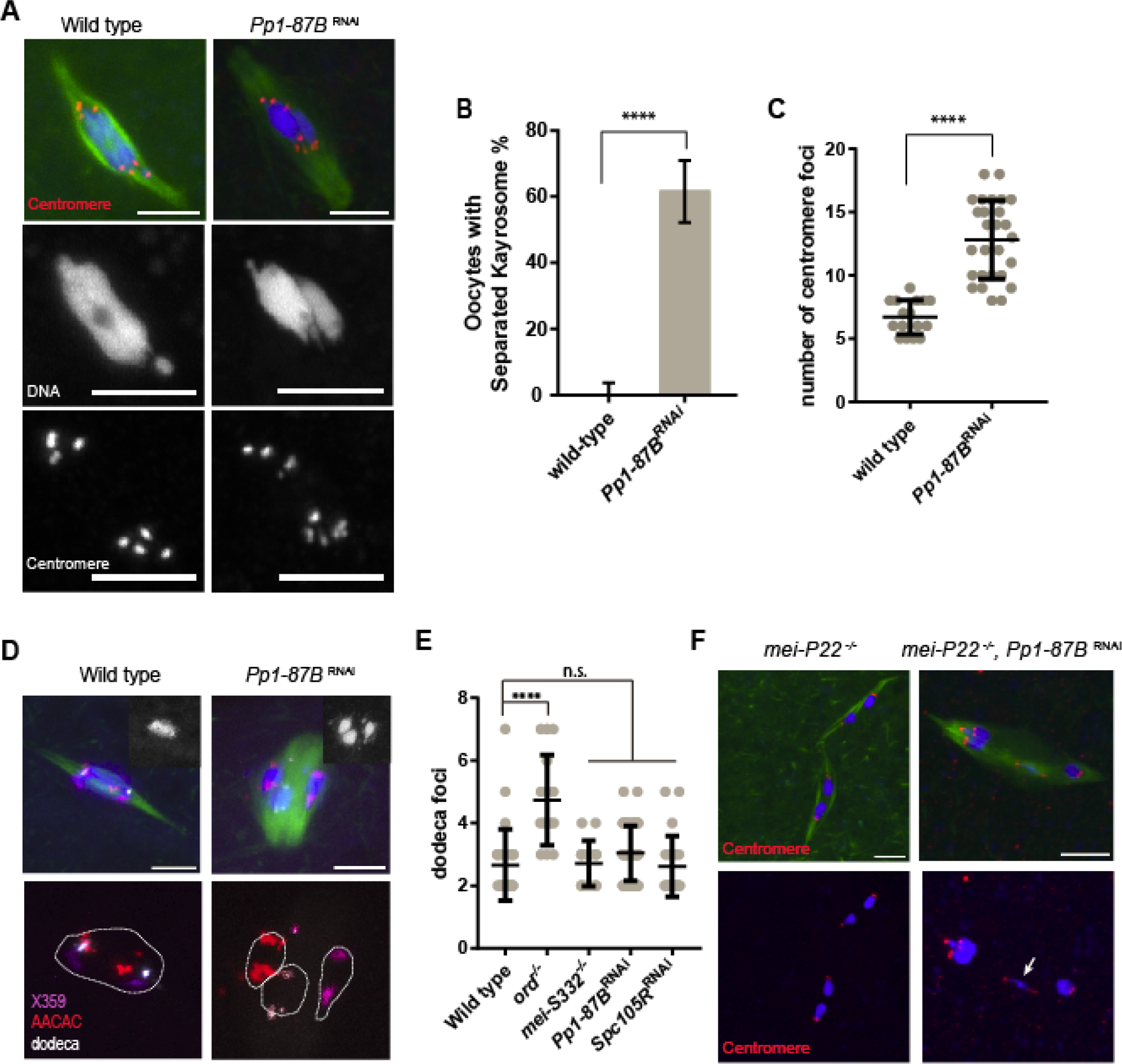
*Pp1-87B* RNAi oocytes have defects in karyosome organization and sister centromere fusion. (A) *Pp1-87B* RNAi oocytes show separated karyosome and sister centromere (red) separation in metaphase I with tubulin in green and DNA in blue. DNA and centromeres are shown in separate channels. In wild-type, the fourth chromosomes sometimes appear as a dot separated from the karyosome. Scale bars indicate 5 μm. (B) Quantification of the separated karyosome phenotype in wild-type (n=20) and *Pp1-87B* RNAi oocytes (n=50). ****= *p* < 0.0001. Error bars indicate 95% confidence interval. (C) Quantification of centromere foci. Error bar shows standard deviation. Number of oocytes: wild type n=16 and *Pp1-87B* RNAi n=27. ****= *p* < 0.0001, (D) Karyosome separation defect in *Pp1-87B* RNAi oocytes. DNA channel is shown in the inset. FISH probes for the X (359 bp repeat, purple), 2^nd^ (AACAC, red) and 3^rd^ chromosome (dodeca, white) were used to detect pericentromeric heterochromatin. The karyosome is outlined in white. Scale bars are 5 μm. (E) Quantification of dodeca foci to detect precocious separation of pericentromeric heterochromatin. Number of oocytes: wild-type (n=27), *ord* (n=15), *mei-S332* (n=14), *Pp1-87B* RNAi (n=50) and *Spc105R* RNAi (n=21). ****= *p* < 0.0001. (F) Recombination defective mutant *mei-P22* displayed homologous chromosome separation indicting precocious anaphase I in oocytes normally arrested in metaphase I. Knocking down *Pp1-87B* in a *mei-P22* mutant background resulted in sister chromatid bi-orientation in meiosis I (arrow).

To determine whether the separate karyosome and centromere separation phenotypes in *Pp1-87B* RNAi oocytes is caused by a general loss of cohesion, we used heterochromatic FISH probes directed to the pericentromeric regions of each autosome to mark homologs. In wild type, two FISH foci are typically observed per homologous chromosome pair, oriented towards opposite poles but within a single karyosome (Figure 1D). If there were loss of arm cohesion, the two homologs could separate and be observed as two FISH foci in separate chromosome masses. We observed that while 62% of *Pp1-87B* RNAi oocytes (n = 50) had separated karyosomes, only 8.5% of the homologs had separated (n =130). These results suggest that arm cohesion is usually retained when PP1-87B is depleted. Hence, the separated karyosome phenotype in *Pp1-87B* RNAi oocytes is due to intact bivalents failing to organize correctly at the center of the spindle.

The same FISH probes could be used to determine if pericentromeric cohesion in *Pp1-87B* RNAi oocytes was affected. We analyzed the number of heterochromatin FISH signals from the dodeca satellite, the most punctate and therefore quantifiable heterochromatic FISH probes. In *ord* mutant oocytes that lack all cohesion, the oocytes had a significantly higher number of dodeca foci (mean = 4.8) compared to wild type (mean = 2.7, Figure 1E). In contrast, the average number of dodeca foci in *Pp1-87B* RNAi oocytes was not significantly higher than wild type (Figure 1E; mean = 3.0), suggesting that pericentromeric cohesion is intact in *Pp1-87B* RNAi oocytes. From these FISH assays we conclude that PP1-87B is only required for maintaining sister centromere cohesion but is dispensable for cohesion of the pericentromeric regions and the chromosome arms in oocytes. To refer to this specific type of cohesion, we will use the term sister centromere fusion.

The defects in *Pp1-87B* RNAi oocytes do result in errors in bi-orientation. In the FISH experiments we observed 5.3% of the homologs in *Pp1-87B* RNAi oocytes were mono-oriented (n=130 v.s. n_wt_=111, *p*=0.016). These results support the conclusion that the sister centromere fusion defect in *Pp1-87B* RNAi oocytes causes problem for homologous chromosomes to bi-orient.

The increased number of centromere foci in *Pp1-87B* RNAi oocytes suggests a defect in sister centromere fusion at meiosis I and hence a lack of co-orientation. When sister centromeres precociously separate dring meiosis I in mouse and yeast, chiasmata can still direct bi-orientation of these homologs, suppressing the consequences of co-orientation defects [9, 13, 25]. Therefore, we used a crossover defective mutant, *mei-P22*, to generate univalents, and knocked down *Pp1-87B* in these oocytes. If the precocious sister centromere separation causes a co-orientation defect, we would expect the univalents in *mei-P22, Pp1-87B* RNAi oocytes can become bi-oriented. Indeed, we observed that 20% of *mei-P22, Pp1-87B* RNAi oocytes had sister chromatids bi-oriented (n = 15, Figure 1F). These results suggest that PP1-87B is required for sister centromere fusion and this facilitate co-orientation in metaphase I of oocytes.

### Co-orientation in Drosophila oocytes requires both cohesin-dependent and cohesin-independent pathways

Both cohesin-dependent and -independent mechanisms of sister centromere fusion have been described. Therefore, we investigated whether loss of sister centromere fusion depends on cohesin release. In addition to PP1-87B, the kinetochore protein SPC105R was also tested because it is the only other protein known to be required for sister centromere fusion in *Drosophila* oocytes [26]. To investigate if cohesin is involved in sister centromere fusion in, we tested if sister centromere separation in *Pp1-87B*- and *Spc105R*-RNAi oocytes depends on known cohesin removal mechanisms by depleting two negative-regulators of cohesin, Wings Apart-like (*wapl*) and Separase (*sse*). If losing a factor required for cohesin removal rescued the sister centromere separation in *Pp1-87B* or *Spc105R* RNAi oocytes, it would suggest the *Drosophila* sister centromere fusion depends on cohesin.

Upon co-expression of *wapl* shRNA with either *Pp1-87B* or *Spc105R* shRNA, the centromeres remained separated (Figure 2A, B), suggesting that Wapl does not regulate sister centromere fusion. While centromere protein localization in *wapl, Spc105R* RNAi oocytes was normal, centromere localization in *wapl*, *Pp1-87B* RNAi oocytes became thread-like instead of punctate (Figure 2A), leading to additional centromere foci when quantified (Figure 2B). The thread-like centromere phenotype suggests that chromosome structure is affected in *wapl*, *Pp1-87B* RNAi oocytes, consistent with previous studies that concluded Wapl was involved in regulating chromosome structure [27, 28].

**Figure 2.**
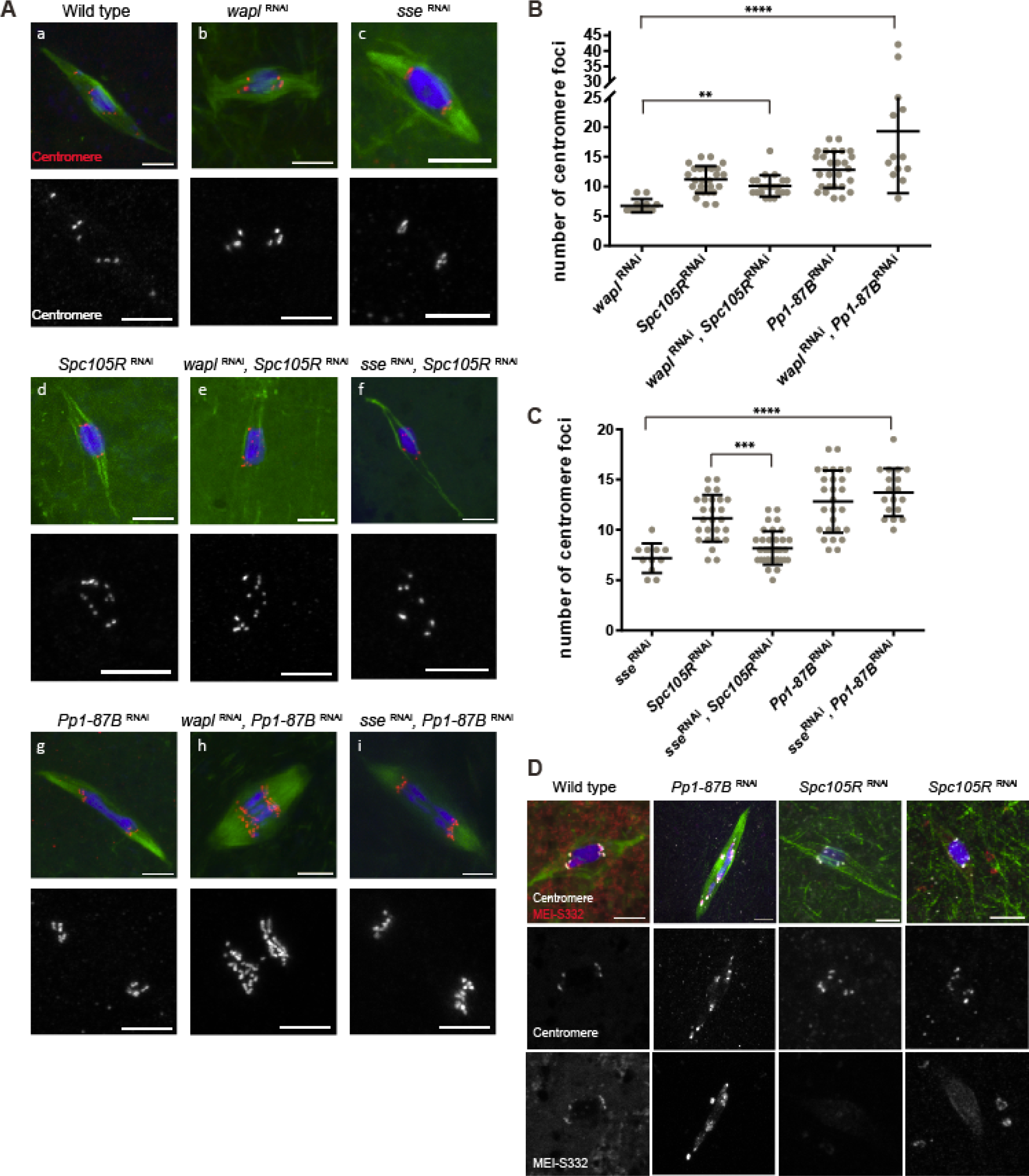
Sister centromere fusion defect rescued by loss of Separase in *Spc105R* RNAi but not *Pp1-87B* RNAi oocytes. (A) Confocal images showing the centromeres (red) in wild-type, *sse* RNAi, and *wapl* RNAi in combination with *Pp1-87B* RNAi or *Spc105R* RNAi. Centromeres are shown in separate channel. Scale bars are 5 μm. (B, C) Dot plots summarize the quantification of centromere foci number in (A). Error bars indicate standard deviation, **= *p* <0.01, ***= *p* <0.001 and ****= *p* <0.0001. Number of oocytes are *wapl* RNAi (12), *Spc105R* RNAi (26), *wapl* RNAi + *Spc105R* RNAi (21), *Pp1-87B* RNAi (27), *wapl* RNAi + *Pp1-87B* RNAi (13), *sse* RNAi (11), *sse* RNAi + *Spc105R* RNAi (36), *sse* RNAi + *Pp1-87B* RNAi (18). (D) MEI-S332 localizes to centromeres and heterochromatin. It has enhanced recruitment to the pericentromeric regions in *Pp1-87B* RNAi oocytes and is decreased in *Spc105R* RNAi oocytes. Two images of *Spc105R* RNAi oocytes show MEI-S332 localization either abolished or greatly reduced. Confocal images are shown with centromeres (white), MEI-S332 (red), tubulin (green) and DNA (blue). Scale bar indicates as 5 μm.

The separated centromere phenotype was rescued in *sse*, *Spc105R* RNAi oocytes (Figure 2A, C; mean = 8.1), suggesting that centromere separation in *Spc105R* RNAi oocytes depends on the loss of cohesins. This is a surprising result because it suggests that Separase is active during meiotic metaphase I [29]. If Separase is active, these results could be explained if SPC105 recruits proteins that protect cohesins from Separase. To test the hypothesis that SPC105R protects cohesins from Separase, we examined the localization of the cohesion protection protein MEI-S332. MEI-S332 localizes to centromere and peri-centromeric regions in wild-type oocytes, as shown by colocalization and substantial non-overlap distribution with the core centromere (Figure S 1). MEI-S332 localization was almost abolished in *Spc105R* RNAi oocytes (Figure 2D). This result suggests that SPC105R is required to recruit proteins that protect cohesins from Separase.

On the other hand, different from the result of *sse*, *Spc105R* RNAi, the separated centromere phenotype was not rescued in *sse*, *Pp1-87B* RNAi oocytes (Figure 2A, C; mean= 13.4). Consistent with cohesin-independence of these phenotypes, the intensity of MEI-S332 in *Pp1-87B* RNAi oocytes was not reduced, and in fact, was increased relative to wild-type (Figure 2D). Aurora B is required for MEI-S332 localization [30], and our results suggest MEI-S332 localization is constrained by antagonism between PP1-87B and Aurora B. These results indicate that sister centromere fusion in *Drosophila* oocytes is regulated through two different mechanisms: the SPC105R pathway that is sensitive to Separase, and the PP1-87B pathway that is Separase independent.

### Separase-independent loss of sister centromere fusion depends on microtubule dynamics

Because the *Pp1-87B* RNAi phenotype was not suppressed by loss of Separase, we investigated cohesin-independent mechanisms for how PP1-87B regulates sister centromere fusion. A critical initial observation was that the spindle volume of *Pp1-87B* RNAi oocytes was larger than wild type (Figure 3A). In addition, PP1-87B was found to localize to the oocyte meiotic spindle (Figure S 2). Based on these observations, we tested the hypothesis that PP1-87B regulates microtubules dynamics by co-depleting proteins known to regulate MT dynamics and KT attachments.

**Figure 3:**
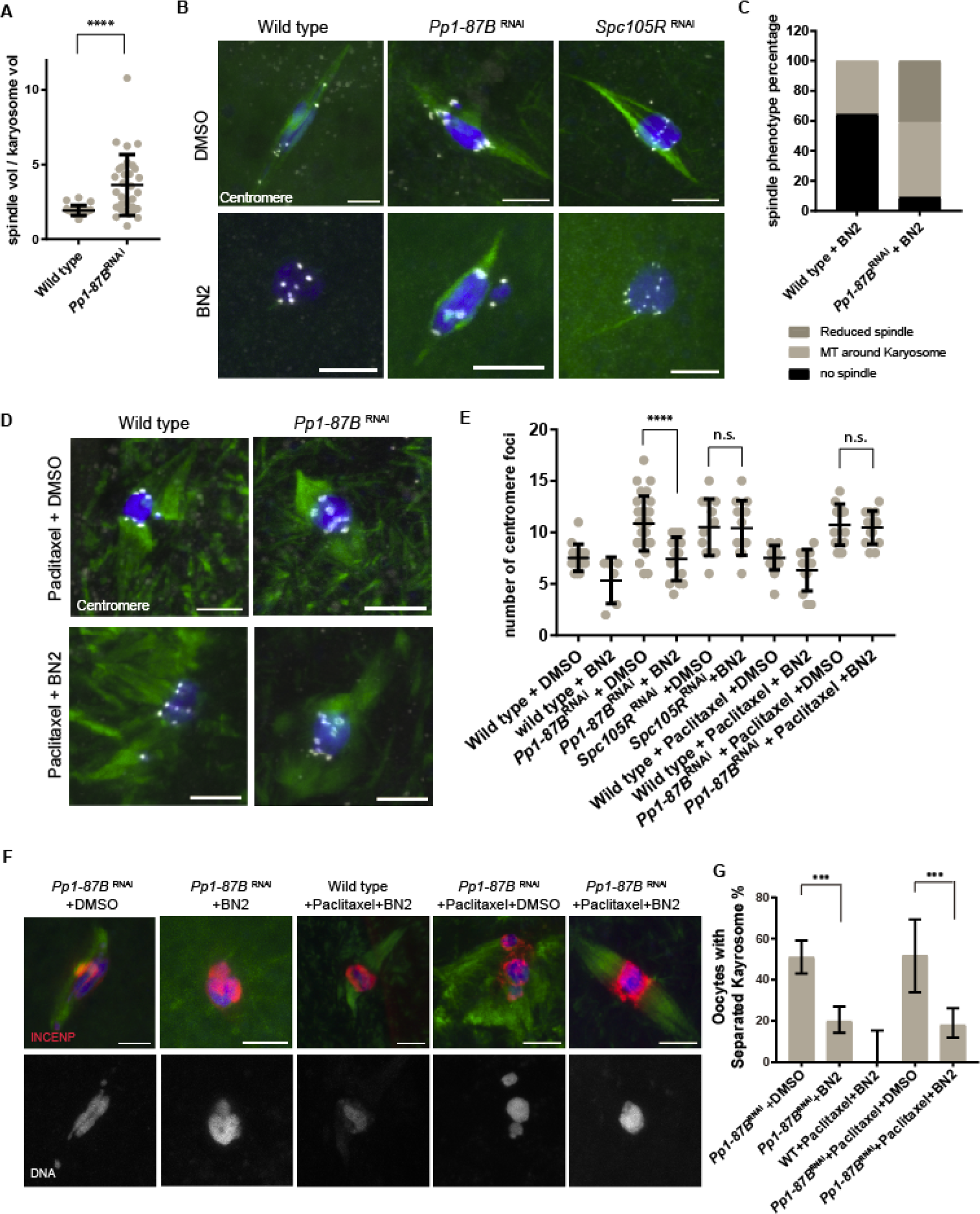
PP1-87B regulation of sister centromere fusion depends on microtubules. (A) Graph showing the spindle volume relative to the karyosome volume. The karyosome volumes remain constant while *Pp1-87B* RNAi oocytes (n=31) had increased spindle volume compared to wild type (22) oocytes. ****= *p* <0.0001 (B) Wild-type, *Pp1-87B* RNAi and *Spc105R* RNAi oocytes treated with 50uM BN2 or the solvent for one hour. All images are shown with DNA (blue), tubulin (green) and centromeres (white), and the scale bars are 5 μm. (C) Quantification of spindle phenotype in wild-type (n=28) and *Pp1-87B* RNAi (n=22) oocytes after one hour of BN2 treatment. (D) Wild type and *Pp1-87B* RNAi oocytes treated with Paclitaxel for 10 minutes followed by either BN2 or DMSO for one hour. Centromeres are marked in white. Scale bars are 5 μm. (E) Quantification of centromere foci in indicated genotypes of oocytes (n= 15, 6, 29, 14, 12, 12, 17, 12, 15 and 13 in the order of the graph). Error bars show standard deviation and ****=p<0.0001. (F) Karyosome organization in *Pp1-87B* RNAi oocytes treated for 10 minutes in Paclitaxel followed by BN2 or DMSO for 30 minutes. INCENP localization is shown in red, DNA in blue and tubulin in green. The single channel of DNA is also shown. Scale bar =5 μm. (G) Quantification from the same experiment in (F). Error bars indicate 95% confidence intervals and ***=p<0.001. The numbers of oocytes were 143,151, 21, 27, 111 in order of the graph.

Aurora B kinase activity is required for spindle assembly in *Drosophila* oocytes [31] and can be antagonized by PP1 in other systems [32]. Furthermore, they have opposite phenotypes: both karyosome and sister centromeres preciously separate in *Pp1-87B* RNAi oocytes but remain together in Aurora B-depleted oocytes [31]. Therefore, we tested whether Aurora B is required for both the karyosome and centromere separation phenotypes of *Pp1-87B* RNAi oocytes. Treatment of oocytes with the Aurora B inhibitor, Binucleine 2 (BN2) [33], caused loss of the spindle (65%, n=29, Figure 3B and C), consistent with previous findings that Aurora B is required for spindle assembly [31]. However, *Pp1-87B* RNAi oocytes showed resistance to BN2 treatment; only 9% had lost the spindle and 50% of oocytes had residual MT around the karyosome (n= 22, Figure 3B and C). Regardless of these residual MTs, the increased number of centromere foci in *Pp1-87B* RNAi oocytes (mean = 13.0) was rescued by BN2 treatment to a level (Figure 3D and E, mean = 7.4) similar to wild-type controls (Figure 3D and E, mean = 7.7).

Similarly, the increased frequency of karyosome separation in *Pp1-87B* RNAi oocytes was rescued by BN2 treatment (Figure 3F and G). In contrast, centromere separation was not rescued by BN2 treatment of *Spc105R* RNAi oocytes (Figure 3B and C, mean = 11.3). These results are concordant with the effects of *sse* RNAi on the *Spc105R* and *Pp1-87B* RNAi phenotypes, and support the conclusion that the maintenance of centromere fusion may occur by at least 2 mechanisms.

Suppression of *Pp1-87B* RNAi oocyte phenotypes by BN2 treatment could have been due to loss of Aurora B activity, or loss of the spindle microtubules. To distinguish between these two possibilities, we treated *Pp1-87B* RNAi oocytes with Paclitaxel to stabilize the spindle prior to BN2 treatment of the oocytes. These oocytes successfully formed spindles; however, 18% showed karyosome separation, a significant decrease compared to the Paclitaxel and solvent-treated RNAi control oocytes and similar to the results from BN2 treatment of *Pp1-87B* RNAi oocytes (Figure 3F and G). This rescue of karyosome separation demonstrates that PP1-87B antagonizes Aurora B in regulating karyosome organization. However, the sister centromeres remained separated in these oocytes (Figure 3D and E, mean = 11.1), suggesting that stabilizing microtubule dynamics in *Pp1-87B* RNAi oocytes can override any effect of inhibiting Aurora B on sister centromere fusion. Based on these observations, we propose that PP1-87B regulates sister centromere separation by regulating microtubules dynamics. However, we cannot rule out the possibility that Aurora B is also required for centromere separation independently of the microtubules.

### Kinetochore-microtubule interactions regulate karyosome organization and sister centromere fusion

The meiotic spindle consists of overlapping microtubules, only a portion of which make contact with kinetochore. To understand which set of microtubules affect PP1-dependent centromere separation and karyosome disorganization, we used knockdowns of kinetochore proteins to specifically abrogate one class of microtubule contracts with the chromosomes. In *Drosophila* oocytes, SPC105R is required for lateral attachments and the localization of NDC80 whereas NDC80 is required for end-on attachments [20]. Thus, we co-depleted PP1-87B and SPC105R (no MT attachments) or NDC80 (lateral MT attachments only) and examined the chromosomes and centromeres. We found that loss of SPC105R, but not NDC80, suppressed the separated karyosome phenotype of *Pp1-87B* RNAi oocytes (Figure 4A and C), suggesting that the separated karyosome phenotype in *Pp1-87B* RNAi oocytes depends on lateral KT-MT interactions. The sister centromeres are already separated in *Spc105R* RNAi oocytes, and co-depletion of both *Pp1-87B* and *Spc105R* did not enhance the effects of either single knockdowns (Figure 4A, B). In contrast, the centromere separation phenotype was rescued in *Ndc80*, *Pp1-87B* double RNAi oocytes (mean = 9.0, Figure 4A, B) but not karyosome disorganization. We conclude that PP1-87B affects karyosome organization through regulating lateral KT-MT attachments and sister centromere fusion through end-on attachments.

**Figure 4.**
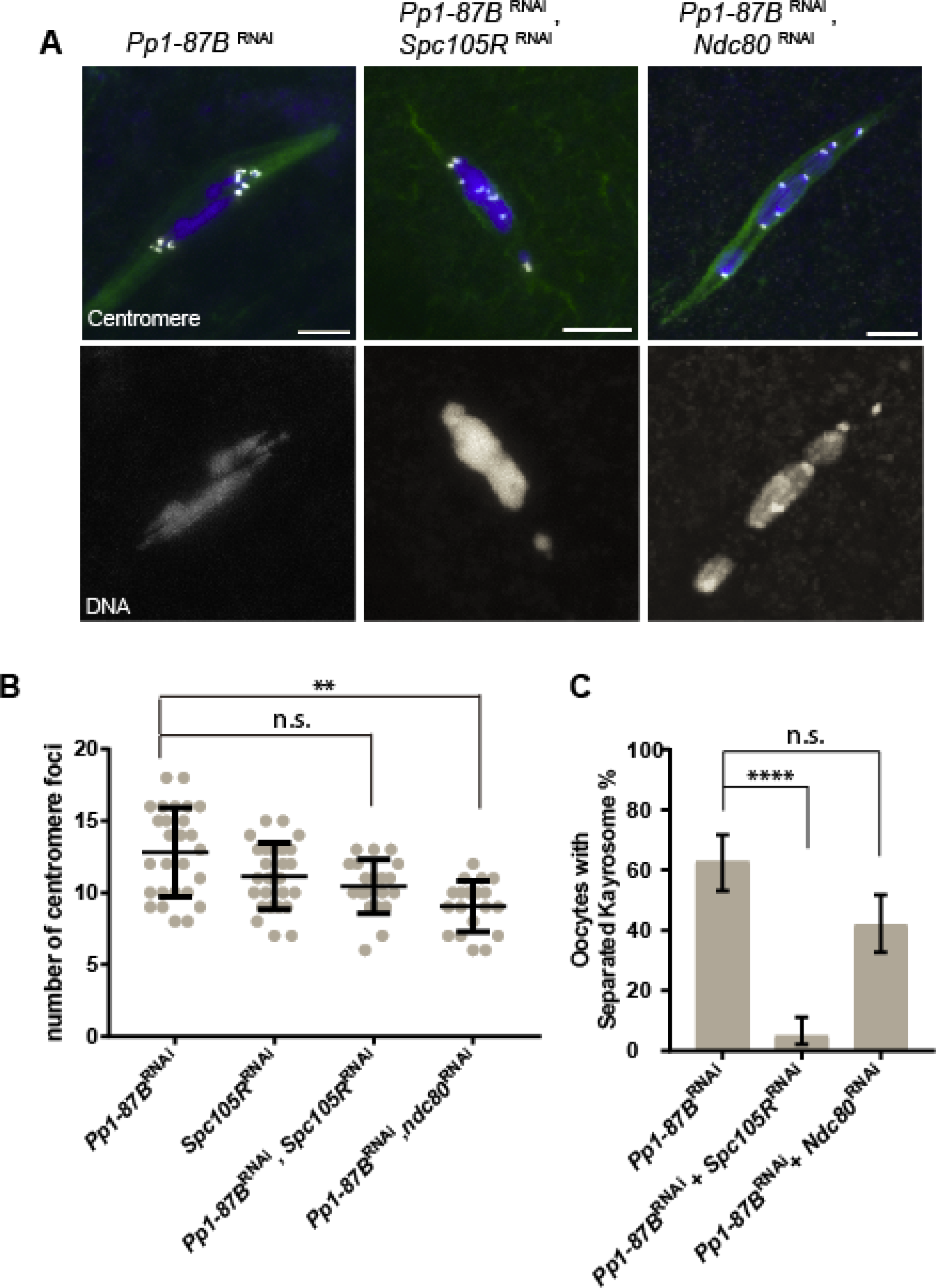
PP1-87B regulates chromosome alignment through lateral attachment and co-orientation via end-on attachment. (A) Confocal images of *Pp1-87B* RNAi oocytes when expressing *Spc105R* RNAi or *Ndc80* RNAi. Centromeres are in white, DNA is shown in blue and tubulin is in green. Single channel image is selected to show DNA in the merge images. Error bars = 5μm. (B) Dot plot shows the number of centromere foci in each genotype. Oocytes numbers are 27, 26, 20 and 19 in order of the graph. Error bars show standard deviation. **=p<0.01. (C) Quantification of oocytes with separated karyosomes. Error bars indicate 95% confidence interval. Numbers of oocytes are 29, 20, and 19 in order of the graph. ****=p<0.0001.

### PP1-87B antagonizes Polo and BubR1 in regulating sister centromere fusion

To identify proteins that function with PP1-87B in regulating end-on KT-MT attachments, we depleted proteins with meiotic functions that are involved in regulating microtubule attachments. Polo kinase localizes to centromeres in *Drosophila* metaphase I oocytes [34] (Figure S 3), and in other organisms has been reported to stabilize KT-MT attachments [35-38]. Unlike Polo in mice [13], *Drosophila polo* RNAi oocytes do not show precocious sister centromere separation at metaphase I [39]. We depleted *polo* with RNAi in either *Spc105R* or *Pp1-87B* RNAi oocytes. Interestingly, we found that centromere separation in both mutant oocytes were rescued by *polo* RNAi (Figure 5A and B, mean = 6.6 and mean = 6.9).

**Figure 5:**
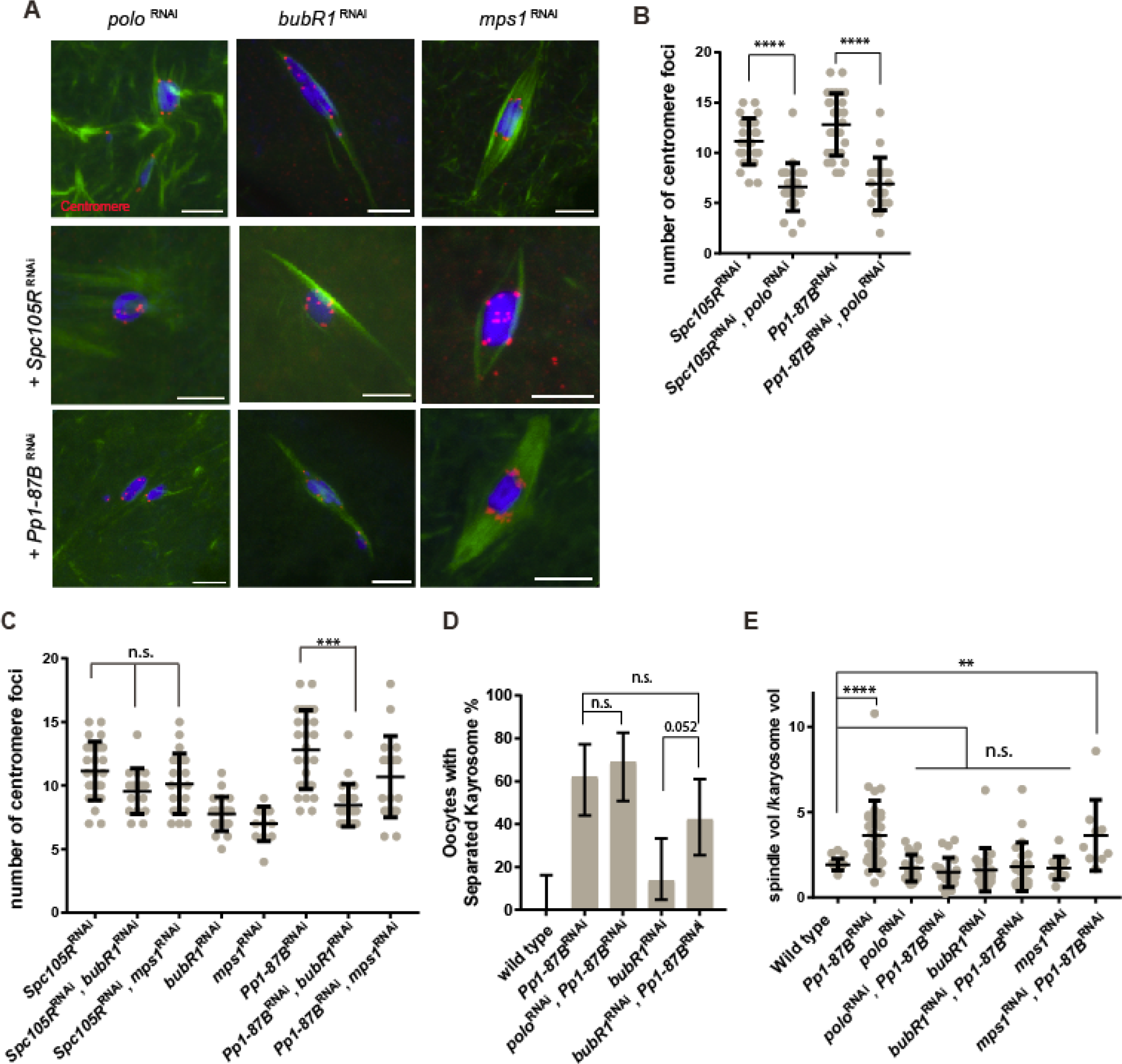
Polo and BubR1 antagonize PP1-87B effects on KT-MT interactions. (A) Confocal images showing *polo, BubR1* or *mps1* RNAi expressed along with *Spc105R* RNAi or *Pp1-87B* RNAi in oocytes. DNA is in blue, tubulin is in green and centromeres are in red. Scale bars = 5μm. (B) Dot plot showing the number of centromeres foci in (A). Oocytes numbers are 26, 24, 27, and 20 in order of the graph. Error bars show standard deviation and ****=p<0.0001. (C) Dot plot showing the number of centromere foci in (A). Oocytes numbers are 26, 16, 21, 22, 12, 27, 22 and 19 in order of the graph. Error bars show standard deviation and ***=p<0.001. (D) Graph showing the percentage of a separated karyosome in oocytes depleted for a variety of kinases in the presence or absence of *Pp1-87B* RNAi. Error bars indicate standard deviation. Numbers of oocytes of each genotype are 20, 31, 21, 19, 22 in order of the graph. (E) Dot plot showing the spindle volume relative to the karyosome volume. Number of oocytes are: 22, 31, 20, 21, 19, 22, 12, 9 in order of the graph. Error bars show standard deviation **= p<0.01 and ****=p<0.0001.

These results indicate that Polo negatively regulates both the cohesion-dependent (through SPC105R) and the microtubule attachment dependent pathways (through PP1-87B) for sister centromere fusion in *Drosophila*.

Two proteins, BubR1 and MPS1, function along with Polo to regulate KT-MT attachments in several organisms [35, 36, 40, 41]. We predicted that depletion of either one could have a similar effect on the *Pp1-87B* oocyte phenotype as *pol*o RNAi. Centromere separation in *Pp1-87B* RNAi oocytes was suppressed by simultaneous knockdown of *BubR1* (Figure 5A, C; mean = 8.5) but not *mps1* (Figure 5A, C; mean = 10.7). A caveat to this negative result is that, based on the non-disjunction rate, MPS1 is only partially depleted in these females (NDJ = 11%, n = 961, compared to a strong *mps1* loss of function mutant, NDJ = 20.2%, n = 231 [42]). Regardless, these results suggest that PP1-87B promotes sister centromere fusion by antagonizing the activities of Polo and BubR1. In contrast, the frequency of oocytes with a separated karyosome phenotype remained similar to *Pp1-87B* RNAi oocytes when PP1-87B were co-depleted with BubR1 (Figure 5D), consistent with the results with NDC80. This result confirms that the separated karyosome phenotype in *Pp1-87B* RNAi oocytes depends on lateral KT-MT interactions.

We propose that PP1-87B inhibits end-on microtubule attachments by antagonizing Polo and BubR1 activities. In support of this conclusion, the increased spindle volume observed in of *Pp1-87B* RNAi oocytes was suppressed by co-depletion of *polo* or *BubR1* (Figure 5E). In summary, several experiments, including drug treatment (Paclitaxel+BN2), depletion of genes that affect KT-MT attachments, and measurements of spindle volume, support the conclusion that PP1-87B regulates KT-MT attachments, and these activities then affect sister-centromere separation and karyosome organization.

### The transverse element of the synaptonemal complex, C(3)G, is required for release of sister centromere fusion

As described above, simultaneous loss of co-orientation and chiasmata can result in bi-orientation of univalent at meiosis I. We observed this phenomenon with codepletion of *PP1-87B* and *mei-P22*. The same experiment was done with *c(3)G*, which encodes a transverse element of the synaptonemal complex (SC) [43], because it is also required for crossing over [44]. Compared to *mei-P22*, however, we got surprisingly different results. First, *c(3)G* mutant females that were depleted of *Pp1-87B* failed to produce mature oocytes. We currently do not know why loss of *c(3)G* and prophase depletion of PP1-87B causes a failure in oocyte development, but it suggests C(3)G has a function in mid-oogenesis after its role in crossing over.

To examine the interaction between C(3)G and PP1-87B, *c(3)G* RNAi was used. To test the efficiency of the *c(3)G* RNAi, *nanos-VP16-GAL4* was used to express the shRNA during early prophase, the frequency of X chromosome non-disjunction (NDJ) was similar to that observed in *c(3)G* null alleles (31%, n = 1647) [44]. In addition, C(3)G localization was absent in the germarium (Figure S 4). These results suggest that this shRNA knockdown recapitulates the null mutant phenotype. For the double depletion we used *mata4-VP16-GAL* that induced shRNA expression later in oogenesis than *nanos-VP16-GAL4*. This was necessary because early expression of *Pp1-87B* shRNA results in a failure to produce oocytes. When using *mata4-VP16-GAL* to express shRNA, C(3)G was present in pachytene, crossing over was not affected (NDJ= 0%, n = 427), but C(3)G was missing from mid-late prophase (Figure 6A, Figure S 4). These results indicate C(3)G is dynamic throughout prophase, and allows us to test if there is a late prophase-metaphase interaction between C(3)G and PP1-87B. Interestingly, RNAi of *c(3)G* rescued the sister centromere separation phenotype in *Pp1-87B*, but not *Spc105R* RNAi oocytes (Figure 6A and B). These results suggest that PP1-87B antagonizes centromeric C(3)G, after most of the SC has been disassembled, to maintain sister centromere fusion at metaphase I. As with other proteins that regulate end-on attachments, the *Pp1-87B* RNAi increased spindle volume phenotype was rescued to wild type levels by co-depletion of *c(3)G* (Figure 6C).

**Figure 6.**
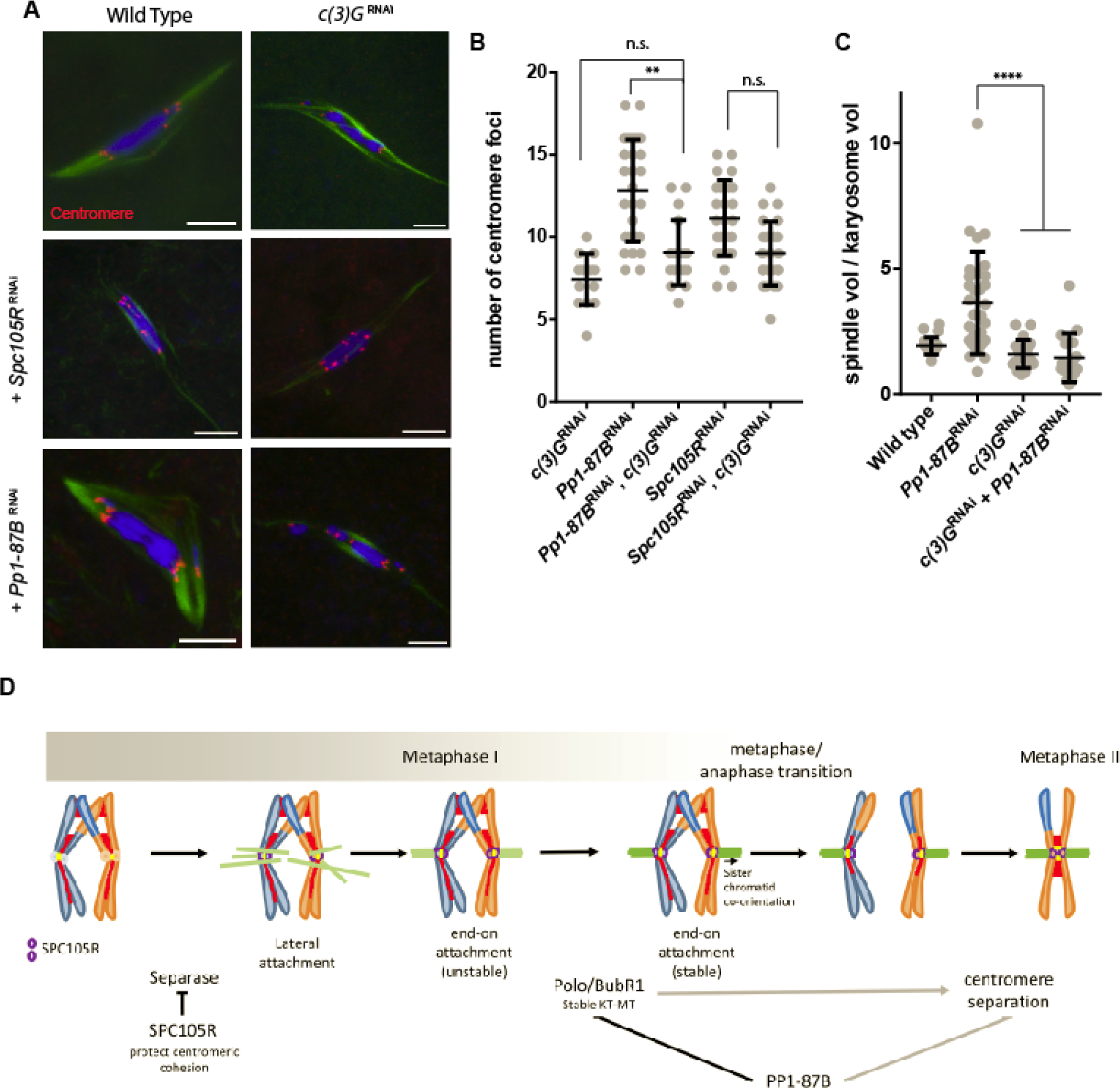
PP1-87B antagonizes C(3)G to regulate sister centromere fusion. (A) Confocal images of oocytes expressing c*(3)G* RNAi in combination with *Pp1-87B* or *Spc105R* RNAi. The centromeres are shown in red in the merged images. Scale bar= 5 μm. (B) Dot plot showing the number of centromere foci in (A). Number of oocytes of each genotype are 14, 27, 18, 26 and 23 in order of the graph. Error bars indicates standard deviation. **=p<0.01. (C) Graph showing the ratio of the spindle volume to the karyosome volume. Number of oocytes are: 22, 31, 20 and 17. Error bars show standard deviation, ****=p<0.0001. (D) Model for regulation of co-orientation in *Drosophila* oocytes.

It is noteworthy that C(3)G is enriched at the centromere regions [45, 46] in pachytene, although its function there is not known. In this location, and because C(3)G has a Polo-binding box, it is possible that C(3)G recruits Polo to the centromere region to regulate microtubule dynamics. However, when examining the localization of Polo in *c(3)G* RNAi oocytes, we did not observe any changes in protein localization compared to wild-type (Figure S 4). Whether C(3)G plays a role in regulating microtubule dynamics through Polo or other independent function to regulate sister centromere fusion needs further investigation.

## Discussion

The fusion of sister centromeres is important for co-orientation in meiosis I, ensuring that sister kinetochores attach to microtubules from the same pole. Release of this attachment must occur early in meiosis II. Based on our results, we propose that the regulation of sister centromere fusion that ensures its release in meiosis II occurs through at least two mechanisms (Figure 6D). First, *Drosophila* centromere fusion depends on loading cohesins at the centromeres that is protected by a kinetochore protein, SPC105R. Second, sister centromere fusion is released in a Separase-independent manner that depends on KT-MT interactions and is inhibited by PP1-87B.

### Sister centromere fusion depends on kinetochore protein SPC105R to protect cohesion from Separase

Assembly of meiosis-specific cohesins at the centromeres probably establishes sister centromere fusion [2]. Indeed, the meiosis-specific cohesin complex SMC1/SMC3/SOLO/SUNN is enriched at *Drosophila* meiotic core centromeres and appears to have this function [14-17]. Depleting Separase in metaphase I *Drosophila* oocytes rescued the precocious centromere separation phenotype caused by loss of SPC105R. Thus, SPC105R protects centromere cohesion, probably by recruiting cohesin protection proteins such as MEI-S332/SGO that subsequently recruit PP2A. A previous finding that *Drosophila Spc105R* mutants enhance defects in Separase function suggest SPC105R may have this function in other cell types [47]. However, Separase activation usually coincides with the entry into anaphase when the APC degrades an inhibitor of Separase, Securin [29]. One possible explanation is that Separase is active prior to anaphase I and cohesion is maintained only by PP2A activity in metaphase I arrested oocytes. This model can explain why knockout of SPC105R in male meiosis does not show a loss of centromere fusion [48]. In male meiosis where there is no cell cycle arrest, Separase may not be active until anaphase, which would make a protective role for SPC105R difficult to observe.

### The transition of sister centromeres from co-orientation to bi-orientation depends on kinetochore-microtubule interactions

Several of our experiments demonstrated that centromere separation in *Pp1-87B* RNAi oocytes was reversible. In addition, treatment of prometaphase I oocytes with the Aurora B inhibitor BN2 showed that centromere separation is reversible in *Pp1-87B* RNAi but not *Spc105R* RNAi. This reversible phenotype is consistent with a mechanism that involves the reorganization of centromere and kinetochore geometry in *Pp1-87B* RNAi oocytes rather than degradation of cohesins. Furthermore, the results from destabilizing microtubule attachments with BN2 treatment suggested that centromere separation in PP1-87B-depleted oocytes depends on KT-MT interactions. In support of this conclusion, we found that PP1-87B affects several spindle-based parameters: it localizes to the meiotic spindle, its knockdown caused an increase in spindle volume, and centromere separation in PP1-87B-deplated oocytes depended on NDC80, Polo and BubR1. These results suggest that stable end-on attachments are required for release of sister centromere fusion. Similar conclusions have been made in *Drosophila* male meiosis. Sister centromere separation in meiosis II does not depend on Separase [49] but does depend on KT-MT interactions [48, 50]. These findings are not limited to *Drosophila*. Classic micro-manipulation experiments in grasshopper cells demonstrated that the switch in meiosis II to separated sister kinetochores requires attachment to the spindle [51]. Based on all these results, we propose that sister centromeres normally separate early in meiosis II by a process that is Separase-independent but microtubule-dependent (Figure 6D).

The mechanism regulated by PP1-87B that regulates KT-MT interactions and maintains sister centromere fusion may be a function utilized in mitotic cells. For example, PP1 has a role in regulating microtubule dynamic in *Xenopus* extracts [52]. In HeLa cells depleted of SDS22, a regulatory subunit of PP1, sister kinetochore distances increase [53], similar to the defect we described here. In budding yeast, suppressing premature formation of stable kinetochore-microtubule attachments is necessary for co-orientation [54]. Finally, our observations are strikingly similar to the phenomenon of cohesin fatigue, where sister chromatids separate in metaphase arrested mitotic cells. Identical to the effect of PP1-87B on centromere separation, cohesion fatigue occurs in a Separase-independent but microtubule-dependent manner [55, 56], however, the mechanism is unknown [57].

Oocytes with a prolonged arrest points, such as metaphase I in *Drosophila*, might prevent cohesion fatigue by concentrating meiotic cohesins at the centromeres and destabilizing KT attachments to reduce MT forces. In *Drosophila* oocytes, the microtubule catastrophe protein Sentin destabilizes end-on KT-MT attachments after the spindle is well established [26]. In fact, active destabilization of kinetochore attachments may be a common feature of oocyte meiosis. Mammalian oocytes also have an extended period of dynamic KT-MT interactions [58], lasting 6-8 hours in mice and up to 16 hours in human [59, 60]. All of these results are in line with our conclusion that oocytes require PP1-87B to prevent premature stable KT-MT attachments and avoiding cohesion fatigue.

### On the role of C(3)G and Polo kinase in cohesion and co-orientation

Depletion of C(3)G suppresses the *Pp1-87B* centromere fusion defect. This result suggests that centromeric SC has a role in negatively regulating sister centromere co-orientation. While the bulk of SC disassembles in late prophase [61, 62], centromeric SC proteins persist beyond pachytene in *Drosophila* and until at least metaphase I in budding yeast and mouse [45, 62-65]. It has also been shown that SC proteins interact with NDC80 complex in two yeast two hybrid experiments [66, 67]. These studies have concluded that centromeric SC is required for bi-orientation of homologs and monopolar attachment. Because both Polo Kinase and C(3)G negatively regulate co-orientation, we hypothesize that C(3)G could be required for Polo Kinase activity, but not localization, at the centromere. Thus, centromeric SC components might be an important mediator of co-orientation.

Co-orientation in yeast and mice depends on Polo kinase, which is recruited by Spo13, Moa1 or Meikin [68]. This is opposite of the known mitotic role of Polo in phosphorylating cohesin subunits and facilitating their removal from binding sister chromatids [69-71]. In yeast meiosis, however, the phosphorylation of cohesin subunits may depend on two different kinases, Casein kinase I and CDC7 [72-74]. Which kinase(s) are required in animals to phosphorylate meiotic cohesins for their removal remains unknown. We have shown that Polo is required for loss of centromeric cohesion, which to our knowledge is the first evidence of its kind in animal meiotic cells.

Unlike mice and yeast, depletion of Polo kinase from *Drosophila* metaphase I oocytes does not cause sister centromere separation [39]. One reason for this difference in Polo function could be that it is required at multiple stages of meiosis and its phenotype may depend on when it is absent. Loss of Polo or BubR1 during early *Drosophila* prophase (pachytene) oocytes leads to loss of SC and cohesion defects [75, 76]. Our experiments deplete Polo after cohesion is established. Alternatively, the function of Polo in co-orientation may not be conserved. Importantly, two features of centromere fusion and co-orientation that are conserved are maintenance depending on SPC105R and separation depending on stabilization of KT-MT attachments. Like SPC105R in *Drosophila*, budding yeast KNL1 is required for meiotic sister centromere fusion and co-orientation and is a target of Polo [18]. The differences between *Drosophila* and mouse or yeast can be explained if SPC105R does not require Polo in order to protect cohesion and the centromeres for co-orientation.

While all previous studies of co-orientation have focused on the establishment of centromere fusion, our results identified several key regulators and provide insights into how sister centromere fusion is maintained in meiosis I and released for meiosis II. In contrast to release of cohesion in most regions of the chromosomes, we propose a Separase-independent mechanism that requires stable kinetochore-microtubule attachments to promotes centromere separation early in meiosis II. While it is well known that regulating microtubule attachments is important for bi-orientation, our result are an example of another reason why KT-MT attachments must be properly regulated to safely navigate the transitions through the two divisions of meiosis.

## Methods

### Drosophila genetics

*Drosophila* were crossed and maintained on standard media at 25°C. Fly stocks were obtained from the Bloomington Stock Center or the Transgenic RNAi Project at Harvard Medical School [TRiP, Boston, MA, USA, flyrnai.org, TRiP, Boston, MA, USA, flyrnai.org, 23]. Information on genetic loci can be obtained from FlyBase [flybase.org, flybase.org, 77] [77].

### RNAi in oocytes: expression and quantification

Most Drosophila lines expressing a short hairpin RNA were designed and made by the Transgenic RNAi Project, Harvard (TRiP) (Table 1). To deplete target mRNA, a cross wss performed to generate females carrying both the *UAS:shRNA* and a *GAL4-VP16* transgene. The shRNA can be induced ubiquitous expression by crossing to *tubP-GAL4-LL7* and testing lethality [78], or *mata4-GAL-VP16* and *osk-GAL4-VP16* for oocyte-specific expression [79]. In this paper, *mata4-GAL-VP16* was primarily used for inducing expression of the *UAS:shRNA* after early pachytene but throughout most stages of oocyte development in the *Drosophila* ovary. This allows for 3-5 days of continuous expression to knockdown the mRNA levels. In some cases, we used the *oskar -GAL4-VP16* transgene [80, 81], which causes a similar knockdown and phenotype in PP1-87B as *mata4-GAL-VP16.* Double RNAi crosses were set up based on the available RNAi lines (Table 2).

**Table 1.**
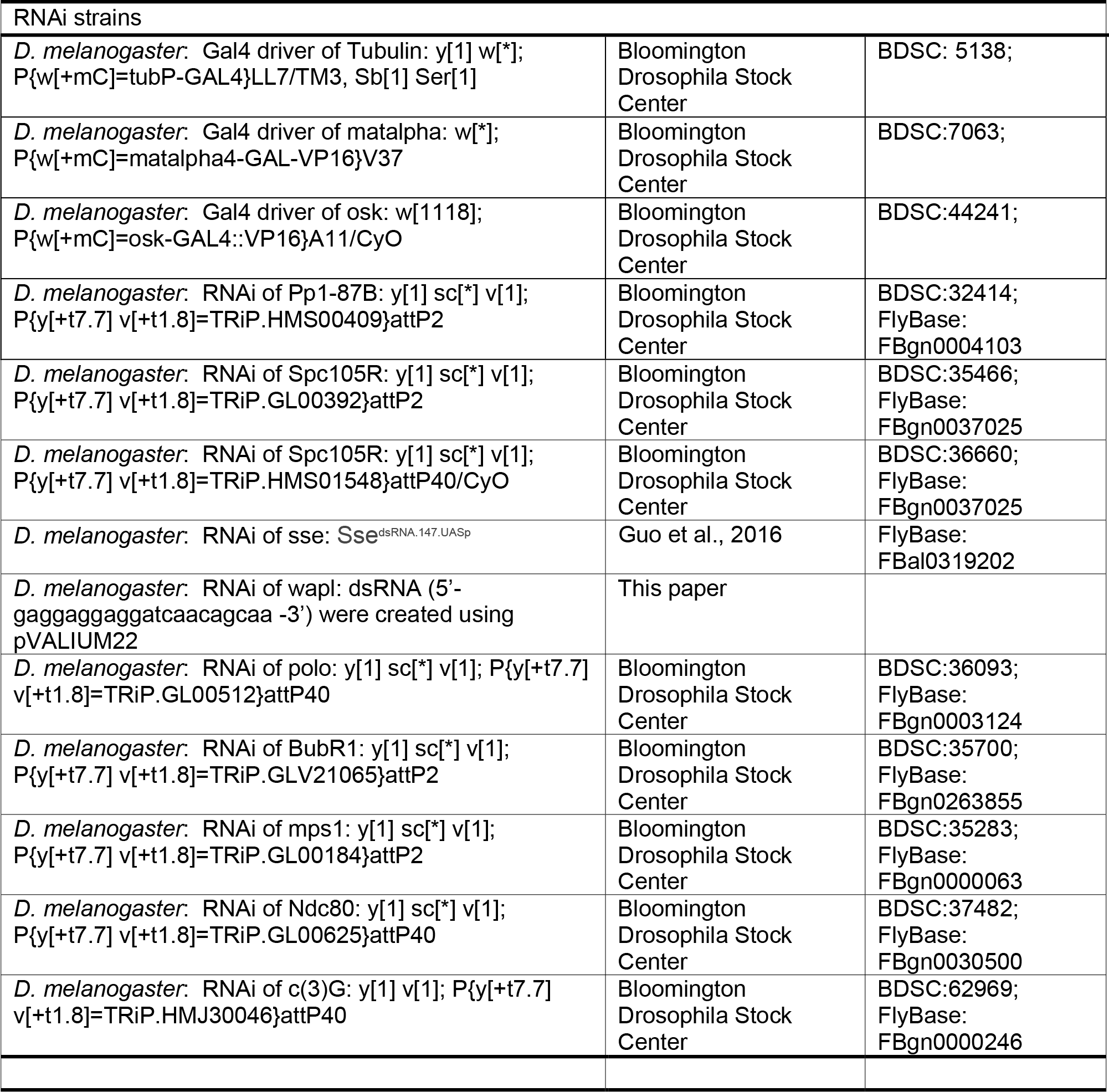

**Table 2:** Transgenes used for Double RNAi

| Driver (GAL) | shRNA line 1 | shRNA line 2 |
| --- | --- | --- |
| mata4-GAL-VP16 | wapl | PP1-87B (HMS00409) or<br>SPC105R (GL00392) |
| mata4-GAL-VP16 | sse<sup>dsRNA.147.UASp</sup> | PP1-87B (HMS00409) or<br>SPC105R (GL00392) |
| mata4-GAL-VP16 | c(3)G (HMJ30046) | PP1-87B (HMS00409) or<br>SPC105R (GL00392) |
| mata4-GAL-VP16 | Polo (GL00512) | PP1-87B (HMS00409) or<br>SPC105R (GL00392) |
| osk-GAL4-VP16 | mps1 (GL00184) | PP1-87B (HMS00409) or<br>SPC105R (GL00392) |
| osk-GAL4-VP16 | bubR1 (GLV21065) | PP1-87B (HMS00409) or<br>SPC105R (GL00392) |
| mata4-GAL-VP16 | Ndc80 (GL00625) | PP1-87B (HMS00409) or<br>SPC105R (GL00392) |
| osk-GAL4-VP16 | SPC105R (GL00392) | PP1-87B (HMS00409) |

For measuring the mRNA knockdown level, total RNA was extracted from late-stage oocytes using TRIzol® Reagent (Life Technologies) and reverse transcribed into cDNA using the High Capacity cDNA Reverse Transcription Kit (Applied Biosystems). The qPCR was performed on a StepOnePlus™ (Life Technologies) real-time PCR system using TaqMan® Gene Expression Assays (Life Technologies), Dm02152292_g1 for *Pp1-87B* and Dm02134593_g1 for the control *RpII140*. Oocyte-specific shRNA expression of *HMS00409* using *mata4-GAL-VP16* resulted in sterility and knockdown of the oocyte mRNA to 35% as measured by RT-qPCR; the same phenotype has been seen when using *osk-GAL4-VP16*, where the mRNA knockdown is also to 35%. For SPC105R, expressing shRNA GL00392 using *osk-GAL-VP16* knocked down the mRNA to 10%.

### Generation of Wapl shRNA lines in *Drosophila*

To generate a *wapl* shRNA line, we followed the protocol in Harvard TRiP center (http://fgr.hms.harvard.edu/trip-plasmid-vector-sets) and targeted *wapl* sequence (5’-gaggaggaggatcaacagcaa -3’) for mRNA knockdown. This 21-nucleotide sequence was cloned into pVALIUM22 and the whole construct was injected into *Drosophila* embryos (*y sc v; attP40*). The mRNA is knocked down to 4% when using *mata4-GAL-VP16* to express the shRNA in oocytes.

### Antibodies and immunofluorescent microscopy

Mature (stage 12-14) oocytes were collected from 100 to 200, 3-4-day old yeast-fed non-virgin females. The procedure is described as in [82]. Oocytes were stained for DNA with Hoechst 33342 (10 μg/ml) and for MTs with mouse anti-α tubulin monoclonal antibody DM1A (1:50), directly conjugated to FITC (Sigma, St. Louis). Additional primary antibodies used were rat anti-Subito antibody [34], rat anti-INCENP [83], guinea pig anti-MEI-S332 [84], rabbit anti-CENP-C [85], rabbit anti-Deterin [86], rabbit anti-Spc105R [87], mouse anti-Polo [88] and rabbit anti-CID (Active Motif). These primary antibodies were combined with either a Cy3, Alex 594 or Cy5 secondary antibody pre-absorbed against a range of mammalian serum proteins (Jackson Immunoresearch, West Grove, PA). FISH probes corresponding to the X359 repeat labeled with Alexa 594, AACAC repeat labeled with Cy3 and the dodeca repeat labeled with Cy5 were obtained from IDT. Oocytes were mounted in SlowFade Gold (Invitrogen). Images were collected on a Leica TCS SP8 confocal microscope with a 63x, NA 1.4 lens. Images are shown as maximum projections of complete image stacks followed by merging of individual channels and cropping in Adobe Photoshop (PS6). CENP-C foci, CID foci, karyosome volume and spindle volume were measured using Imaris image analysis software (Bitplane) and graphs were plotted and statistics calculated using Graphpad Prism software.

### Binuclein 2 treatment assay

To inhibit Aurora B, oocytes were incubated with either 0.1% DMSO or 50 μM BN2 in 0.1% DMSO for 60 minutes prior to fixation in Robb’s media. To stabilize MTs, oocytes were incubated with either 0.1% DMSO or 10 μM Paclitaxel (Sigma) in 0.1% DMSO for 10 minutes, followed by 50 μM BN2 plus 10 μM Paclitaxel in 0.1% DMSO for 60 minutes.

### Quantification and statistical analysis

Statistical tests were performed using GraphPad Prism software. All the numbers of the centromere foci or spindle/karyosome volume were pooled together and ran one-way ANOVA followed by post hoc pairwise Tukey’s multiple comparison test. Details of statistical evaluations and the numbers of samples are provided in the figure legends.

Figure S 1: **MEI-S332 localization does not co-localize with centromere**. Related to Figures 1 and 2. Representitive picture of wild type oocytes staining MEI-S332 (red) and CID (white) is shown and measured the intensity of flourensent. MEI-S332 localizes to both the pericentromeric and centromeric regions. Scale bar is 5 μm.

Figure S 2: **Localization of PP1-87B to meiotic spindle**. Related to Figures 1 and 4. An epitope-tagged version of PP1-87B was expressed from a UASP transgene using *mata4-GAL-VP16.* HA-PP1-87B is in red, tubulin in green and DNA in blue and scale bars are 5 μm.

Figure S 3: **Polo localization does not change in *c(3)G* RNAi oocytes but decreases in *Spc105R* RNAi oocytes.**Related to Figures 3 and 5. Wild-type, *c(3)G* RNAi, and *Spc105R* RNAi oocytes with DNA in blue, tubulin in green, Polo in red and CID in white. Single channels are shown in white. All images are maximum projections and scale bars are 5 μm.

Figure S 4: **C(3)G is knockdown by shRNA expressed in the germline. (**A) C(3)G (red) forms thread-like structure in the germarium, and retains them in oocytes of stages 2-5 of the vitellarium. (B) When *nanos-VP16-GAL4* expressed *c(3)G* shRNA in early prophase, C(3)G expression was abolished. (C) When *mata4-GAL-VP16* expressed *c(3)G* shRNA, C(3)G localization was present in germarium early pachytene, but absent in the stages 2-5 of the vitellarium. Scale bars are 10 μm.

